# Old Compound, New Context: Naphthocyclinones Mediate Microbial Competition in Wild Bee Nests

**DOI:** 10.64898/2026.10.24.761495

**Authors:** Roman Makitrynskyy, Imen Nouioui, Marlen Jando, Yuliia Kovalova, Jolantha Swiderski, Cathrin Spröer, Boyke Bunk, Andrey Yurkov, Yvonne Mast

## Abstract

Although thousands of microbial specialized metabolites have been identified, the ecological functions of most of these compounds remain unknown. Naphthocyclinones are aromatic polyketide antibiotics that were discovered more than five decades ago, yet their biological role has remained elusive. During investigation of the microbial community associated with the oligolectic ivy bee (*Colletes hederae*), a diverse bacterial community that included several *Streptomyces* species and members of the class *Bacilli* was recovered from fermented nest provisions. One *Streptomyces* isolate, B-004^T^ (DSM 121525^T^ = LMG 34598^T^), was identified as representing a novel species based on polyphasic taxonomic studies, for which the name *Streptomyces hederae* sp. nov. is proposed. This strain showed potent antibacterial activity against co-isolated Gram-positive bacteria as well as *Paenibacillus larvae*, the causative agent of American foulbrood. Bioassay-guided fractionation coupled with liquid chromatography-mass spectrometry analysis led to the identification of naphthocyclinone congeners as the active metabolites. Whole-genome sequencing and genome mining revealed a type II polyketide synthase biosynthetic gene cluster closely related to that of the previously characterized naphthocyclinone pathway. Targeted deletion of the cluster-encoded ketosynthase gene (*B-004_07360*) abolished naphthocyclinone production and resulted in the complete loss of antibacterial activity, whereas heterologous overexpression of the *Streptomyces* antibiotic regulatory protein gene *papR2* significantly increased antibiotic production confirming the function of the naphthocyclinone biosynthetic gene cluster. Furthermore, we found that naphthocyclinone biosynthesis is induced by the presence of sympatric bacterial isolates from bee nests. Together, our findings provide the first evidence for an ecological function of naphthocyclinones as mediators of bacterial competition in wild bee nests and highlight their potential as antimicrobial substances against *P. larvae*.

## Introduction

Solitary bees play an important role in maintaining terrestrial biodiversity and supporting agricultural ecosystems through their pollination services (1,2). Unlike social honeybees (*Apis mellifera*), which rely on complex hive structures and social immunity, solitary bees face unique environmental challenges during their reproductive cycles (3). The ivy bee, *Colletes hederae*, is an oligolectic species that is strictly specialized on ivy pollen and constructs underground nests, and provisions its brood cells with a mixture of pollen and nectar (4). These fermented nest provisions represent a nutrient-rich microenvironment that is potentially highly susceptible to microbial spoilage, while ground-built nests are additionally exposed to infiltration by pathogens. Under such conditions, solitary bees may benefit from associations with microorganisms that protect larval provisions from microbial invasion (5). In line with this, a recent study on the oligolectic solitary bee *Andrena vaga* demonstrated that microbial communities associated with ground nests and larval provisions are likely to provide essential functions to this bee (6). Specifically, two yeast species are involved in the detoxification of bee provisions by reducing the salicin content from pollen, while another species dominates on eggs and may provide protection to the nest.

Several important pathogens of honey and wild bees belong to the spore-forming class *Bacilli* (7). Most notably, *Paenibacillus larvae* is the causative agent of American foulbrood, a severe infectious disease of honeybees capable of decimating larval populations and destroying entire colonies (8). Although the mechanisms of pathogenesis and disease control have been extensively studied in honeybees, the ecological interactions between sympatric microorganisms and potential pathogens within the nests of solitary bees such as *C. hederae* remain poorly understood (5). A better understanding of the mechanisms underlying microbial suppression of opportunistic pathogens in these natural microhabitats may facilitate the discovery of novel antimicrobial compounds and their ecological functions.

Actinomycetes, particularly members of the genus *Streptomyces*, are renowned producers of structurally diverse secondary metabolites and account for the majority of clinically important antibiotics (9). Their genomes encode numerous biosynthetic gene clusters (BGCs), many of which remain silent or are only poorly expressed under laboratory conditions (10). To overcome the transcriptional silence and low expression of BGCs, several activation strategies have been developed. One such strategy is the heterologous overexpression of genes encoding *Streptomyces* antibiotic regulatory proteins (SARPs), which has proven particularly effective for activating silent BGCs and increasing metabolite production (11,12).

Streptomycetes are soil-dwelling microorganisms, some of which are known to establish mutualistic associations with insects, where antibiotic production protects the host or its food resources from microbial pathogens (9). A well-studied example is the defensive mutualism between fungus-growing ants and antibiotic-producing actinomycetes, in which antimicrobial metabolites protect the fungal cultivars against specialized pathogens, such as *Escovopsis* (13,14). Similar defensive roles have been described in solitary wasps, where symbiotic streptomycetes produce antibiotic cocktails containing compounds such as piericidins, streptochlorin, and nigericin that protect developing wasp larvae (15,16). In termite nests, nest-associated streptomycetes contribute to colony hygiene by suppressing microbial pathogens and enhancing host survival, demonstrating that antibiotic-mediated defense can remain stable over long-term host–microbe associations (17,18). Moreover, evolutionary analyses indicate that antibiotic-based defense in such symbioses can be remarkably stable over long timescales (19). Despite the ecological importance of actinomycetes in insect-associated microbiomes, the natural functions of most of the secondary metabolites they produce remain poorly understood. Naphthocyclinones are asymmetric dimeric aromatic polyketides of the benzoisochromanequinone family, belonging to the type II polyketides (20,21). They were originally isolated more than five decades ago from *Streptomyces arenae.* Several stereochemical variants have been described, of which the β- and γ-forms exhibit strong antibacterial activity against Gram-positive bacteria (22). Although their chemical structures and biosynthetic pathway have been extensively characterized (23), little is known about their ecological function or the environmental contexts in which they are naturally produced. (6)Expanding the search for beneficial microbial associates of oligolectic bees, the present study investigated microorganisms isolated from the fermented nest provisions of the wild bee *C. hederae*. Here, we describe the identification of a *Streptomyces* strain producing naphthocyclinones, which display antibacterial activity against co-isolated sympatric *Bacilli* and the pathogen *P. larvae*. By combining genome sequencing, bioactivity assays, and targeted mutagenesis, we functionally linked the observed antibacterial activity to a naphthocyclinone BGC. We further demonstrated that heterologous overexpression of the SARP-type regulatory gene *papR2* significantly enhanced and accelerated naphthocyclinone production. Our findings provide the first experimental evidence for an ecological function of naphthocyclinones in mediating bacterial competition within bee-associated microbial communities and show that these long-known aromatic polyketides contribute to the suppression of a major bee pathogen.

## MATERIALS AND METHODS

### Bacterial strains, plasmids, and cultivation conditions

Plasmids and bacterial strains used in this study are listed in Table S1. All *Escherichia coli* strains were grown in Luria–Bertani (LB) and 2×YT media at 37°C, supplemented with appropriate antibiotics when required. *Streptomyces* strains were cultivated on soya flour mannitol agar (SFM), R5 and R5A, R2YE agar, tryptone soya broth (TSB) (24), NL19 (20 g soya flour, 20 g mannitol, 1 L tap water, pH 7.3), GYM (4 g glucose, 4 g yeast extract, 10 g malt extract, 1 L distilled water, pH 7.2), S (4 g peptone, 4 g yeast extract, 4 g K2HPO4, 2 g KH2PO4, 10 g glycine, 800 mL distilled water, after sterilization 10 g glucose and 0.5 g MgSO4 in 200 mL distilled water), and oatmeal agar (OAT; 36 g oatmeal, 20 g agar, 1 L tap water, pH adjusted to 7.3) media at 28°C. For growth in liquid cultures, strains were incubated in baffled flasks on a rotary shaker at 180 rpm. Co-isolated strains were grown on tryptone soya agar (TSA), J-agar (DSM medium 262), and Columbia blood agar (DSM medium 429) at 28°C. Columbia blood agar and J-agar were used to cultivate *P. larvae* DSM 7030^T^ at 28°C under microaerophilic conditions. Apramycin (50 µg mL⁻¹), carbenicillin (100 µg mL⁻¹), chloramphenicol (25 µg mL⁻¹), hygromycin B (100 µg mL⁻¹), and kanamycin (50 µg mL⁻¹) were added to the growth media when required.

### Procedures for DNA manipulation

The basic procedures for DNA manipulation were performed as described by Sambrook et al. (25) for *E. coli* and by Kieser et al. (24) for *Streptomyces*. Oligonucleotides used in this study are listed in Table S2. All enzymes were purchased from New England Biolabs (NEB). Polymerase chain reactions (PCRs) were performed using Q5 High-Fidelity DNA Polymerase (NEB). All constructs were verified by Sanger sequencing performed at Eurofins Genomics (Eurofins Genomics Europe Shared Services GmbH; Anzinger Str. 7a; 85560 Ebersberg; Germany) using custom primers and by PCR and/or restriction analysis.

### Sample collection and microbial isolation

Samples of *C. hederae* nests were collected by Dr. Hanna Gardein (Julius Kühn-Institut, Braunschweig, Germany) on 16 October 2024 in the Westliches Ringgebiet district of Braunschweig, Lower Saxony, Germany (GPS: 52.2675086, 10.5016338). Approximately 20 mg of fermented nest provision obtained from a nest was used for the isolation procedure. Samples were thoroughly resuspended in 1 mL of sterile 0.9% (wt/vol) NaCl solution by vortexing. Serial dilutions (10⁻²–10⁻^3^) were prepared from the suspension, and 100 µL aliquots were plated onto OAT agar plates. Plates were incubated at 28°C and analyzed after 10 days.

Individual colonies were streaked onto OAT and TSA plates and incubated until sufficient growth was observed. Single-colony isolates were subsequently cryopreserved as glycerol stocks at −80°C. Following strain identification and characterization, selected isolates were deposited in the DSMZ culture collection under the respective accession numbers (Table 1).

**Table 1.**
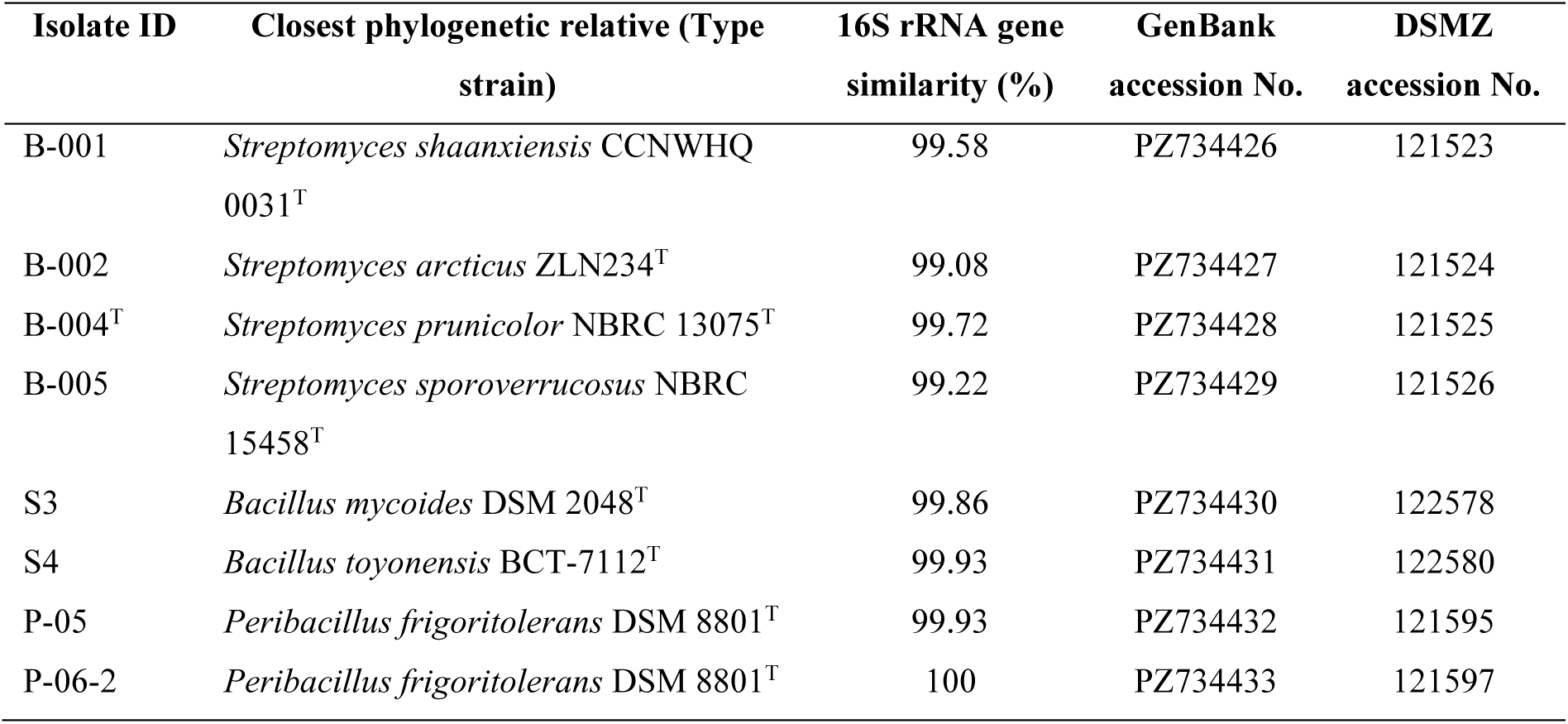
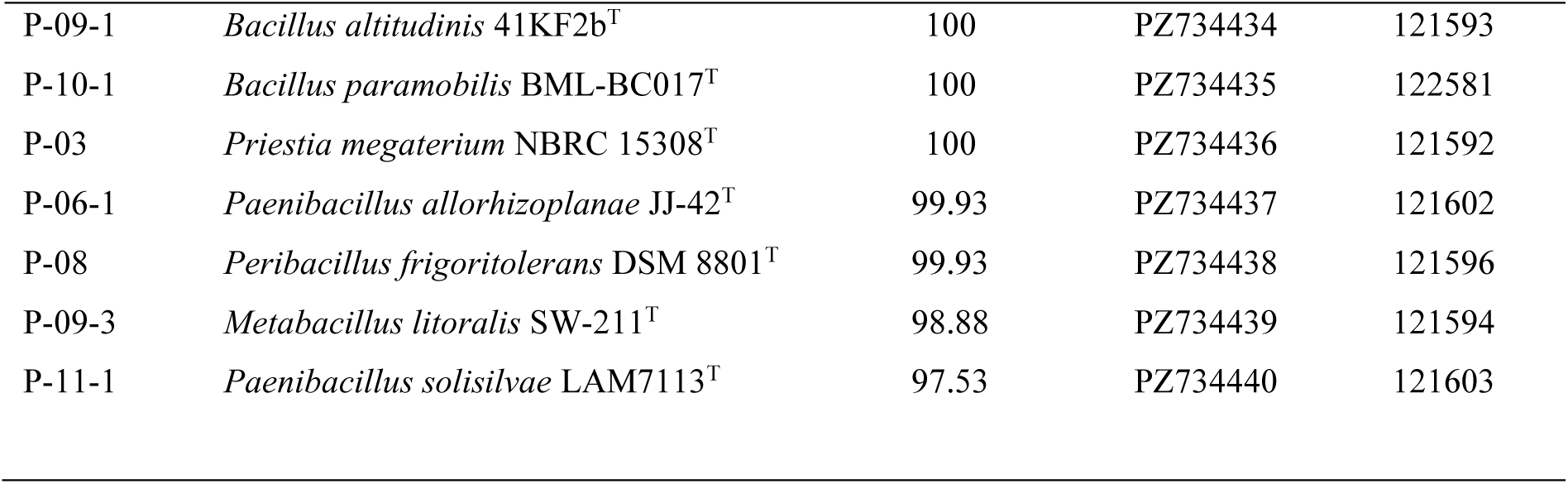
Taxonomic identification and accession numbers of bacterial isolates from the *C. hederae* fermented provision.

### Strain identification and phylogenetic analysis

Genomic DNA was extracted from actively growing cultures using the NucleoSpin Microbial DNA Kit (Macherey-Nagel, Düren, Germany) according to the manufacturer’s instructions. The 16S rRNA gene was amplified by PCR and sequenced by the Sanger method at Eurofins Genomics (Ebersberg, Germany). Nearly complete 16S rRNA gene sequences (>1,400 bp) were obtained, deposited in GenBank under respective accession numbers (Table 1), and compared with the 16S rRNA gene sequences of type strains of validly published species available in the EZBioCloud server (https://www.ezbiocloud.net) (26). Pairwise 16S rRNA gene sequence similarities between the isolates and their closest phylogenetic neighbors were determined using the EZBioCloud server. *Streptomyces hokutonensis* DSM 102214^T^ was identified as the closest phylogenetic relative of strain B-004^T^ based on the 16S rRNA gene sequence analysis and was therefore included as a reference strain in the comparative phenotypic and taxogenomic analyses. DSM 102214^T^ was obtained from the DSMZ culture collection. A neighbor-joining (NJ) phylogenetic tree based on the 16S rRNA gene sequence was constructed using MEGA version 12 with 1,000 bootstrap replications (27–30).

### Genome sequencing, assembly, and taxogenomic analyses

High-quality genomic DNA was extracted from actively growing cultures using the MasterPure Gram-positive DNA Purification Kit (Lucigen, Middleton, WI, USA) according to the manufacturer’s instructions. Whole-genome sequencing was performed on the PacBio Vega platform. Raw HiFi reads were assembled *de novo* using the PacBio microbial genome assembly pipeline. The 16S rRNA gene sequence extracted from the genome was aligned with the PCR-derived 16S rRNA gene sequence using Basic Local Alignment Search Tool (BLASTN) (31) (https://blast.ncbi.nlm.nih.gov/Blast.cgi). The two sequences were identical, confirming the authenticity of strain B-004^T^. Genome quality was assessed based on assembly statistics, and the assembled genome sequence was deposited in GenBank under accession number (ID:1522304, SUB16454956). Phylogenomic and comparative genomic analyses, including digital DNA-DNA hybridization (dDDH) were performed using the Type (strain) genome server (TYGS) (https://tygs.dsmz.de/) (32,33). The dDDH values were calculated using the recommended formula *d*4 implemented in TYGS (32). Average nucleotide identity (ANIm) values between the strain B-004^T^ and its closest phylogenomic relatives were calculated using JSpecies WS server (34,35). Growth properties of B-004^T^ and related biochemical, enzymatic, and chemotaxonomic analysis are described in Supplemental Text S1.

### Protoplast transformation

Transformation of *Streptomyces* sp. B-004^T^ was performed by polyethylene glycol (PEG)- mediated protoplast transformation according to the protocol described by Kieser et al. (24), with minor modifications. Briefly, the strain was grown in S medium for 48 h at 28°C and 180 rpm. Protoplasts were generated by lysozyme treatment (1 mg mL⁻¹), mixed with 1 µg plasmid DNA, and transformed by PEG-mediated DNA uptake. Following regeneration on R2YE agar for 18 h, plates were overlaid with soft nutrient agar containing the appropriate antibiotics, and transformants were incubated at 28°C until colonies appeared.

### Generation of an in-frame ketosynthase deletion mutant

An in-frame deletion of the ketosynthase gene *B-004_07360,* was generated by homologous recombination to generate a double crossover event. A 7.64-kb DNA fragment containing *B-004_07360* (1,266 bp) and approximately 3.2-kb up- and downstream homologous regions was amplified by PCR from genomic DNA of strain B-004^T^ using the primer pair RM-b-KA-syn_long_f and RM-b-KA-syn_long_r (Table S2). The resulting amplicon was cloned into EcoRV-digested pBluescript, yielding the plasmid pBl-B-004-KA-syn_long. An in-frame deletion of *B-004_07360* was generated by PCR-based mutagenesis, as described previously (36). Primers RM-b-KA-PCRmut_f and RM-b-KA-PCRmut_r (Table S2) were used to amplify a 9.5-kb DNA fragment lacking *B-004_07360*, using pBl-B-004-KA-syn_long as the template. The amplicon was phosphorylated using T4 polynucleotide kinase and ligated with a 1-kb apramycin resistance cassette, *aac*(*3*)*IV*, PCR-amplified from plasmid pLEERECJ (37) using primers RM_aac(3)IV_pLee_f and RM_aac(3)IV_pLee_r, generating pBl-B-004-ΔKA::aac(3)IV. A 7.46-kb DNA fragment containing *aac*(*3*)*IV* in place of gene *B-004_07360* and flanked by homologous regions was amplified by PCR from pBl-B-004-ΔKA::aac(3)IV using primers RM-b-KA-syn_long_f and RM-b-KA-syn_long_r. The fragment was subcloned into the hygromycin resistance (*hyg*)-bearing suicide vector pKGLP2 (38), digested with EcoRV, generating pKGLP2-B-004-ΔKA::aac(3)IV. The final construct was introduced into *Streptomyces* sp. B-004^T^ via PEG-mediated protoplast transformation, followed by screening for apramycin-resistant and hygromycin-sensitive colonies, indicative of a double-crossover event and plasmid loss. In total, nine independent clones of *Streptomyces* sp. B-004^T^ Δ*07360::aac*(*3*)*IV* were obtained, and replacement of gene *B-004_07360* was confirmed by PCR using specific primer sets as described in Fig. S1.

### Antibacterial activity assays

Bioactivity was assessed using disk diffusion assays against a panel of reference microorganisms, including Gram-positive bacteria (*Kocuria rhizophila* DSM 348, *Paenibacillus larvae* DSM 7030ᵀ), the Gram-negative bacterium (*Escherichia coli* Δ*tolC* JW5503–1), yeasts (*Saccharomyces cerevisiae* DSM 70449^T^, *Lodderomyces* (formerly *Candida*) *parapsilosis* DSM 5784^T^, *Solicoccozyma aeria* DSM 120404, *Metschnikowia pulcherrima* DSM 121680, *Metschnikowia reukaufii* DSM 121684), and the filamentous fungus *Aspergillus brasiliensis* DSM 1988, as well as bacterial co-isolates obtained together with the *Streptomyces* strains from the fermented provision of *C. hederae*.

*Streptomyces* strains were initially cultivated in 25 mL of TSB to generate precultures. After 48 h, 1 mL of preculture was used to inoculate 50 mL of R5, GYM, and NL19 media, followed by incubation for 4 days at 28°C in 500-mL Erlenmeyer flasks on a rotary shaker (180 rpm). For preliminary bioactivity screening, 20 mL of fermentation broth was centrifuged at 5,000 rpm, and the supernatant was concentrated in vacuo to a final volume of 4 mL. Aliquots (50 µL) of the concentrated supernatants were applied onto sterile paper disks, which were then placed onto double-layer agar plates. The bottom layer consisted of TSA or J-agar containing 2% (wt/vol) agar, while the upper layer consisted of the same medium supplemented with the test organism in soft agar (0.7% wt/vol agar). Plates were incubated at 4°C for 1 h to allow diffusion and then at 28°C for 16–20 h (48 h for *Paenibacillus* strains). Bioassays were performed at least triplicates.

### Overexpression of the SARP regulatory gene *papR2*

To overexpress *papR2* in *Streptomyces* sp. B-004^T^, the ϕBT1-derived integrative plasmid pDS300, which carries the hygromycin resistance gene *hyg* and drives *papR2* transcription under the control of the constitutive *ermE\** promoter was used (12). Both, pDS300 and the corresponding empty vector pIJ10257 (39) (used as a negative control), were individually introduced into *Streptomyces* sp. B-004 via protoplast transformation. Individual hygromycin-resistant colonies were selected after 6–8 days of incubation on R2YE agar. For naphthocyclinone production, the resulting strains *Streptomyces* sp. B-004/pDS300 and *Streptomyces* sp. B-004/pIJ10257, were cultivated in R5 medium for 6 days as described above.

### Metabolite extraction and chemical analysis

S*treptomyces* sp. B-004 was cultivated in R5 medium for naphthocyclinone production as described above. Fermentation broth (20 mL) was extracted with an equal volume of ethyl acetate or n-butanol for 1 h at room temperature under constant vertical rotation. After centrifugation at 5000 rpm for 10 min, the organic phase was dried using a centrifugal evaporator (SP Genevac EZ-2, “Low BP” program). The extracts were dissolved in 800 µL of methanol and filtered through 0.2 µm PVDF-filters. Next, the samples were directly used for LC-MS analysis. In parallel, 15 µL of extracts were used in antibacterial activity assays.

Low-resolution LC–MS data were acquired using an Agilent 1260 HPLC system coupled to a G6125B mass spectrometer, equipped with a reversed-phase Poroshell C18 column (Agilent; 2.7 µm particle size, 100 × 3 mm). Separation was performed using water as mobile phase A and acetonitrile as mobile phase B, both containing 0.1% (vol/vol) formic acid as a solvent modifier. Elution was carried out at a flow rate of 0.5 mL min⁻¹ using the following gradient program: 0 min, 95% A; 0.5 min, 95% A; 18.5 min, 5% A; 20.5 min, 5% A; 20.8 min, 95% A; 25 min, 95% A. Full-scan mass spectra were collected over an m/z range of 150–2000 in both positive and negative electrospray ionization (ESI) modes. The following source parameters were used: capillary voltage, 3000 V; nebulizer gas pressure, 35 psi; drying gas flow rate (N₂), 10 L min⁻¹; drying gas temperature, 350 °C.

High-resolution LC–MS (HRMS) analysis was performed using an Agilent 1290 HPLC system coupled to an Agilent Accurate-Mass Quadrupole Time-of-Flight (Q-TOF) mass spectrometer. Analytes were separated on an ACQUITY BEH C18 column (Waters; 1.7 µm particle size, 100 × 2.1 mm) using water containing 0.1% (vol/vol) formic acid as mobile phase A and acetonitrile containing 0.1% (vol/vol) formic acid as mobile phase B. Elution was performed at a flow rate of 0.55 mL min⁻¹ using the following gradient program: 0 min, 95% A; 20 min, 0% A; 21 min, 0% A; 31 min, 95% A. Data were acquired in positive and negative electrospray ionization (ESI) modes over an m/z range of 200–2000.

Semi-preparative HPLC analysis was performed using an Agilent 1260 HPLC system coupled to an Agilent automatic fraction collector (G1364F). Separation was carried out on a reversed-phase C18 column (Zorbax SB, 9.4 × 150 mm, 5 µm particle size; Agilent Technologies) using water containing 0.1% (vol/vol) formic acid as mobile phase A and acetonitrile containing 0.1% (vol/vol) formic acid as mobile phase B. The following elution gradient was used: 0 min, 95% A; 0.5 min, 95% A; 13.5 min, 5% A; 16.5 min, 5% A; 17 min, 95% A; 20 min, 95% A. The mobile phase was delivered at a flow rate of 2.5 mL min⁻¹. Fractions were collected every minute during the run, and the solvents were evaporated using a centrifugal evaporator (SP Genevac EZ-2, “HPLC” program). Dry residues were reconstituted in methanol and used directly for bioassays or LC-MS analysis.

### Analysis of biosynthetic gene clusters

Putative BGCs in the genome of *Streptomyces* sp. B-004 were identified using antiSMASH v8.0 (40), followed by manual annotation. BLAST analysis (31) was performed to screen for orthologous genes in the GenBank database. Biosynthetic gene cluster comparisons and visualization were performed using clinker (41).

## Results

### Isolation of a diverse bacterial community from *C. hederae* nest provisions

Cultivation of fermented nest provisions of *C. hederae* on media selective for bacterial isolation yielded a total of 18 bacterial strains. Since the primary focus of this study was on the preservation of bee provisions, we specifically searched for actinomycetes, which are prolific producers of specialized metabolites. Isolates displaying an aerial mycelium morphology typical of streptomycetes were prioritized for further characterization. This screening yielded seven actinobacteria isolates, all of which were assigned to the genus *Streptomyces* based on 16S rRNA gene sequence analysis. Dereplication revealed that three isolates likely represented clonal duplicates. Comparison of the remaining four 16S rRNA gene sequences with those of validly published type strains using the EZBioCloud server (26) identified their closest phylogenetic relatives, sharing sequence similarities ranging from 99.08% to 99.72% (Table 1). The remaining 11 isolates belonged to the class *Bacilli* and were assigned to the genera *Bacillus*, *Peribacillus*, *Priestia*, *Paenibacillus*, and *Metabacillus*. 16S rRNA gene sequence similarities to the closest related type strains ranged from 97.53% to 100% (Table 1). All microorganisms isolated in this study were deposited in the DSMZ culture collection under the accession numbers (Table 1).

### *Streptomyces* sp. B-004T displays potent antibacterial activity against co-occurring *Bacilli*

To evaluate the biosynthetic capacity of the isolated streptomycetes, the strains were cultivated in NL19, R5 and GYM fermentation media, and concentrated supernatants from 5-day-old fermentation broths were screened against a panel of test organisms. This panel comprised the Gram-positive bacterium *K. rhizophila* DSM 348, the Gram-negative *E. coli* Δ*tolC*, as well as several yeast species, including baker’s yeast *S. cerevisiae* DSM 70449^T^, the bee-associated *M. pulcherrima* DSM 121680 and *M. reukaufii* DSM 121684, the opportunistic human pathogen *L. parapsilosis* DSM 5784^T^, and the soil-borne *S. aeria* DSM 120404. Additionally, the filamentous fungus strain *A. brasiliensis* DSM 1988 was included in the bioassay. Antimicrobial activity was detected exclusively in the supernatant samples of strain B-004^T^ cultured in NL19 and R5 media against the Gram-positive test strain *K. rhizophila* (Fig. S2A). To assess whether the observed antibacterial activity also affected sympatric microorganisms, the co-isolated members of the class *Bacilli* were included in subsequent bioassays. Consistent with the inhibition observed for the Gram-positive *K. rhizophila*, all co-isolated *Bacilli* exhibited sensitivity to the supernatants of strain B-004^T^ (Fig. S2A). Because members of the genus *Paenibacillus* were recovered among the co-isolated *Bacilli*, the inhibitory activity of strain B-004^T^ was subsequently evaluated also against the type strain of *P. larvae* DSM 7030ᵀ, the causative agent of American foulbrood. Indeed, both the culture supernatant and organic solvent extracts strongly inhibited the growth of the type strain DSM 7030ᵀ (Fig. 1). Finally, to determine whether antimicrobial metabolites were also present in the natural habitat, fermented nest provisions were sequentially extracted with water and methanol and tested against *K. rhizophila*. Only the methanolic extract showed low growth inhibition, thereby confirming the presence of antimicrobial compounds directly within the bee nest environment (Fig. S2B).

**Fig. 1.**
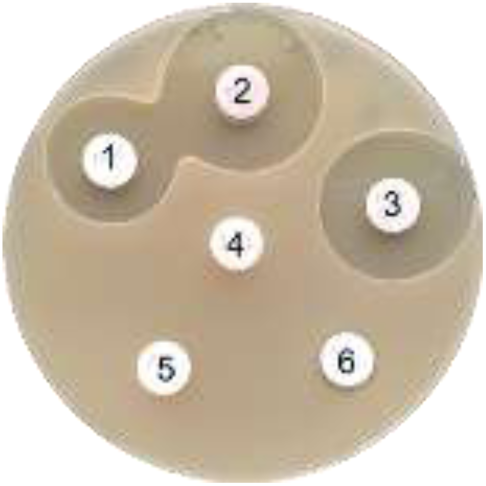
Growth inhibition of *P. larvae* DSM 7030 by *Streptomyces* sp. B-004^T^ grown in NL19. Discs correspond to: 1, concentrated supernatant; 2, n-butanol extract; 3, ethyl acetate extract; 4, concentrated NL19 medium control; 5, ethyl acetate extract of NL19 medium; and 6, n-bu-tanol extract of NL19 medium.

### Naphthocyclinones account for the antibacterial activity of strain *Streptomyces* sp. B-004T

To identify the metabolites responsible for the observed antimicrobial activity, *Streptomyces* sp. B-004^T^ was cultivated in NL19 medium, and the ethyl acetate culture extract was fractionated via semi-preparative HPLC. The fractions were dried *in vacuo*, reconstituted in methanol, and then tested for antibacterial bioactivity against *K. rhizophila*. The activity was detected in fractions eluting between 12 and 15 min (Fig. 2A).

**Fig. 2.**
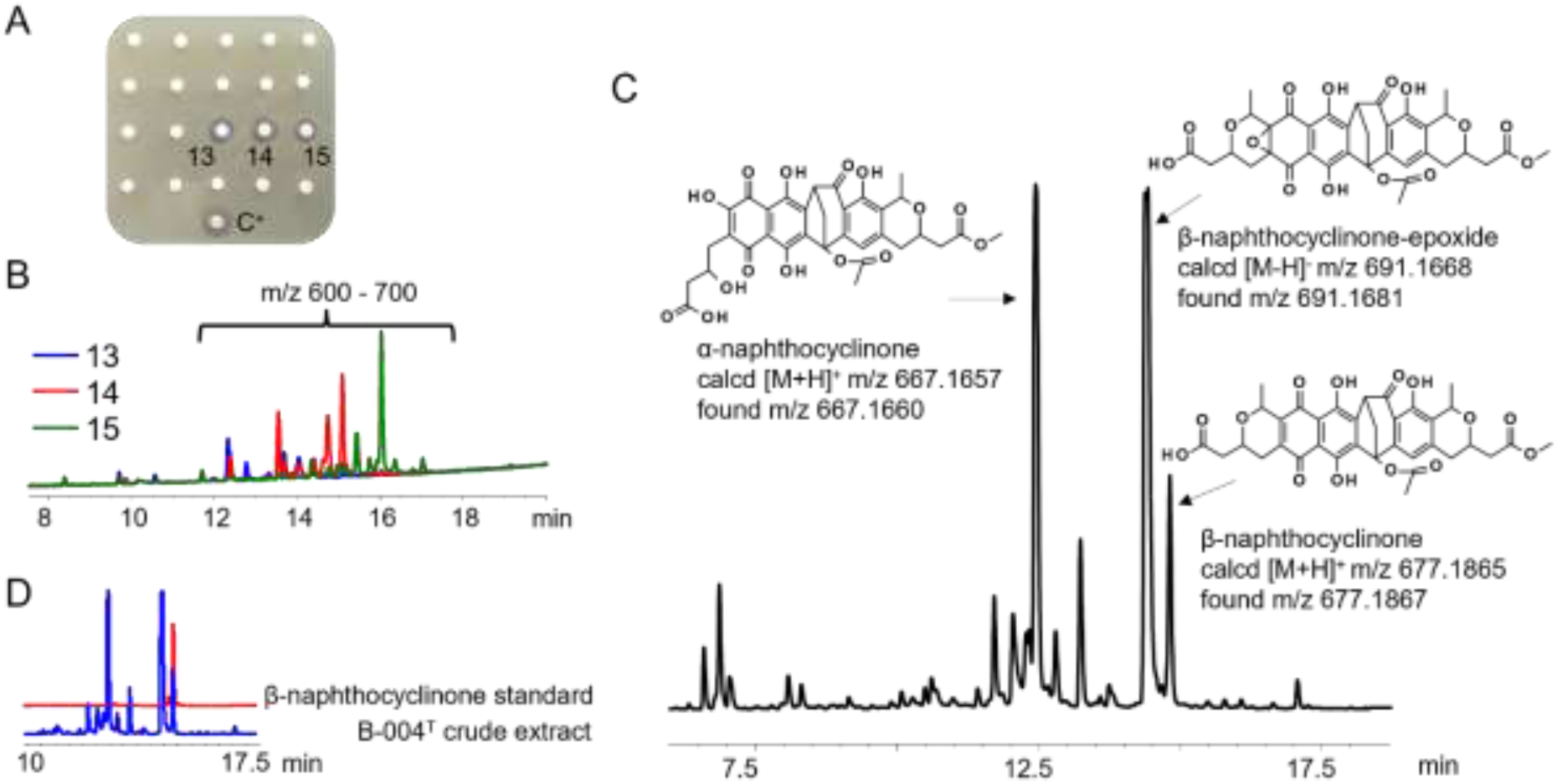
Bioactivity-guided purification and LC-MS/HRMS structural identification of naphthocyclinone congeners from *Streptomyces* sp. B-004^T^. (**A**) Antimicrobial activity assay of semi-preparative HPLC fractions against *K. rhizophila*. Discs 13, 14, and 15 represent collected fractions showing clear zones of growth inhibition; C⁺ – ethyl acetate crude extract of B-004^T^. (**B**) Overlaid HPLC traces of active fractions 13 (blue), 14 (red), and 15 (green) tracked at 254 nm. (**C**) HRMS base peak chromatogram of the *Streptomyces* sp. B-004^T^ crude extract, annotated with chemical structures and high-accuracy mass analysis data. (D) Overlaid HPLC traces recorded at 254 nm comparing the retention time and chromatographic profile of the commercially available authentic standard of β-naphthocyclinone (red trace) against the metabolites of the *Streptomyces* sp. B-004^T^ crude ethyl acetate extract (blue trace).

LC-MS analysis of the bioactive fractions revealed several overlapping peaks (Fig. 2B), which were further analyzed by high-resolution mass spectrometry (HRMS). This analysis yielded a number of protonated molecular ions within the *m/z* range of 600-700 [M+H]^+^. The most abundant mass peaks were detected at *m/z* 677.1867 [M+H]^+^, 667.1660 [M+H]^+^, and 691.1681 [M-H]^-^. Dereplication against the Dictionary of Natural Products (42,43) revealed that these molecular ions closely matched β-naphthocyclinone, α-naphthocyclinone, and β-naphthocyclinone-epoxide, respectively (Fig. 2C and S3). In addition, the UV-Vis spectra (Fig. S3) of the target peaks closely correlated with previously published literature data for different naphthocyclinone congeners (20,22,44). The identity of β-naphthocyclinone was confirmed by LC-MS analysis of a commercially available authentic standard of β-naphthocyclinone, which exhibited an identical retention time as the corresponding metabolite detected in the culture extract of *Streptomyces* sp. B-004^T^ (Fig. 2D).

### Genome mining identifies the naphthocyclinone biosynthetic gene cluster

To link the naphthocyclinone production to the cognate BGC, the *Streptomyces* sp. B-004^T^ genomic DNA was sequenced using the PacBio Vega platform. The resulting high-quality genome assembly comprised 11.19 Mb and consisted of two contigs, representing an 11,177,889 bp chromosome and a 9,048 bp plasmid. The GC content of the chromosome was 70.0%, whereas the plasmid exhibited a slightly lower GC content of 67.8%.

To identify BGCs encoding specialized metabolites, the genome sequence of *Streptomyces* sp. B-004^T^ was analyzed using antiSMASH v. 8.0. A total of 36 putative BGCs was predicted, highlighting the specialized metabolic potential of strain B-004^T^. These included clusters for nonribosomal peptide synthetases (NRPS), terpenes, lanthipeptides (class I), butyrolactones, siderophores, ribosomally synthesized and post-translationally modified peptides (RiPPs), different classes of polyketide synthases (PKS), and various hybrid BGCs (Fig. S4).

Among the predicted BGCs, a type II polyketide synthase (PKS) cluster showed high sequence similarity to the previously characterized α-naphthocyclinoic acid/fogacin BGC from *Streptomyces arenae* (21). Detailed sequence analysis of region 1.26 revealed a highly conserved core region encoding the minimal PKS complex, including orthologs of the ketosynthase α (*ncnA*), ketosynthase β/chain-length factor (*ncnB*), and acyl carrier protein (*ncnC*). In addition, the BGC contained genes encoding cyclases, oxygenases, and tailoring enzymes, including the cofactor-independent oxidase NcnN (Fig. 3), previously shown to be involved in the oxidative dimerization of fogacin monomers and the formation of naphthocyclinone congeners (23), as detected by our LC-MS analysis.

**Fig. 3.**
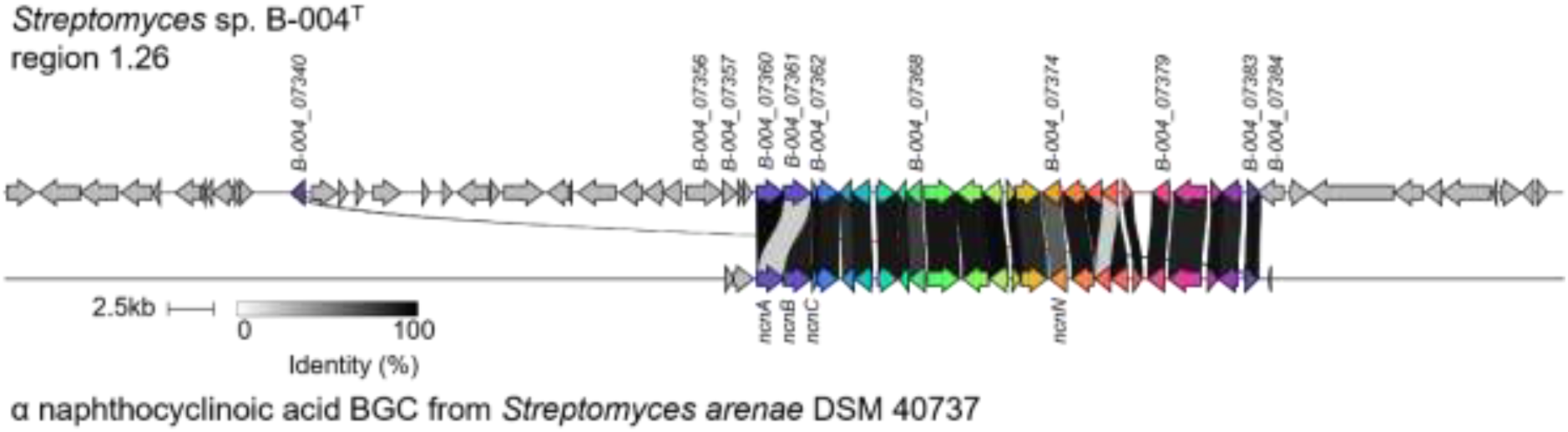
Comparative synteny and genetic architecture of the naphthocyclinone BGCs in *Streptomyces* sp. B-004^T^ and *Streptomyces arenae* DSM 40737. Linear alignment generated using clinker showing the conserved genomic synteny between *Streptomyces* sp. B-004^T^ region 1.26 and the characterized α-naphthocyclinoic acid BGC from *Streptomyces arenae* DSM 40737 (GenBank accession: JAKQYH010000014.1). Core minimal PKS genes (*ncnA*, *ncnB*, *ncnC*), the gene encoding the key cofactor-independent oxidase involved in naphthocyclinone dimerization (*ncnN*), and genes encoding putative transcriptional regulators are indicated.

### Targeted mutagenesis confirms the ecological role of naphthocyclinones

To validate the functionality of the identified naphthocyclinone BGC and confirm its role in antibacterial activity against the test organisms, the core biosynthetic ketosynthase gene *B-004_07360* was deleted by homologous recombination as shown in Fig. S1. The coding sequence of *B-004_07360* was replaced in-frame by the apramycin resistance cassette *aac*(*3*)*IV*, yielding the mutant strain *Streptomyces* sp. B-004^T^ Δ*07360::aac*(*3*)*IV*. Correct gene replacement was verified by PCR analysis (Fig. S1).

Deletion of *B-004_07360* markedly altered the phenotypical appearance of the strain. When grown on R5 agar, the *Streptomyces* sp. B-004^T^ Δ*07360::aac*(*3*)*IV* mutant did not produce the characteristic reddish-brown pigmentation, in contrast to the parental strain (Fig. 4A), suggesting impaired naphthocyclinone biosynthesis of the mutant. To analyze the impact of the gene deletion on metabolite production, the B-004^T^ wild-type (WT) and mutant strains were grown in parallel in R5 medium for five days. LC-MS analysis of culture ethyl acetate extracts revealed the complete absence of naphthocyclinone-related peaks in the Δ*07360::*aac(*3*)*IV* mutant, whereas the characteristic naphthocyclinone peaks were readily detected in the WT sample (Fig. 4B). This result confirmed that the predicted BGC, including the core biosynthetic gene *B-004_07360*, is essential for naphthocyclinone biosynthesis.

**Fig. 4.**
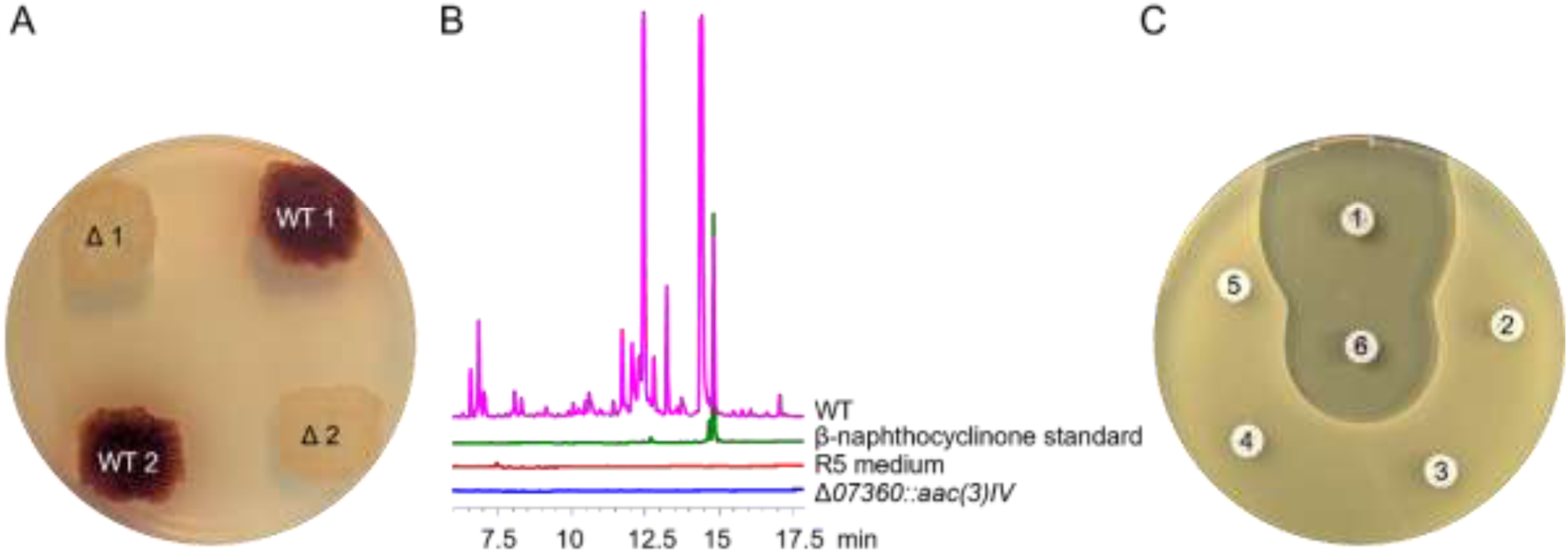
Validation of the ecological role of naphthocyclinones via targeted gene deletion in *Streptomyces* sp. B-004^T^. (**A**) Phenotypes of the wild-type *Streptomyces* sp. B-004^T^ (WT 1, WT 2) and the minimal PKS ketosynthase mutant (Δ1, Δ2) cultivated on R5A for 5 days. (**B**) Overlaid HPLC-UV chromatograms (λ = 254 nm) of ethyl acetate culture extracts from the *Streptomyces* sp. B-004^T^ strains. The wild-type profile (WT, magenta trace) displays prominent naphthocyclinones peaks, including β-naphthocyclinone, which aligns with the commercial standard (green trace). Naphthocyclinone production is entirely abolished in *Streptomyces* sp. B-004^T^ Δ*07360::aac*(*3*)*IV*, (blue trace). Red trace is the extract from uninoculated R5 medium. (**C**) Disc diffusion assay against *P. larvae* DSM 7030^T^ confirming the loss of antimicrobial activity in the mutant. Paper discs saturated with ethyl acetate culture extracts from: 1, wild-type B-004^T^; 2, R5 medium; 3 and 4, B-004^T^ Δ*07360::aac*(*3*)*IV* mutant clone 1 and 2, respectively; 5, methanol; 6, commercial β-naphthocyclinone standard reference (20 mcg).

To determine the impact of *B-004_07360* deletion on antibacterial activity, ethyl acetate extracts of the Δ*07360::*aac(*3*)*IV* mutant were tested against the panel of sensitive Gram-positive bacteria described above. Unlike the wild-type extracts, the mutant extracts showed no antibacterial activity against any of the co-isolated *Bacillus* strains or *P. larvae* (Fig. 4C and S5). These findings demonstrate that naphthocyclinones are solely responsible for the antibacterial activity of strain B-004^T^ against co-isolated Gram-positive bacteria and *P. larvae*.

### SARP overexpression enhances naphthocyclinone production

Analysis of the naphthocyclinone region in *Streptomyces* sp. B-004^T^ revealed several putative regulatory genes, including genes encoding a stand-alone LuxR-family transcriptional regulator (*B-004_07340*), a LuxR-family regulator and its cognate sensor histidine kinase (*B-004_07383* and *B-004_07384*, respectively), a two-component regulatory system (*B-004_07356* and *B-004_07357*), a TetR-family transcriptional regulator (*B-004_07368*), as well as a regulator belonging to the SARP family (*B-004_07379*) (Fig. 3). Members of the SARP-family regulators are well-known direct transcriptional activators of numerous BGCs in actinomycetes (11). Because heterologous expression of the SARP regulator *papR2*, an essential activator of pristinamycin biosynthesis in *Streptomyces pristinaespiralis* (45), has previously been shown to activate or enhance the production of diverse specialized metabolites (11,12), its potential to activate naphthocyclinone biosynthesis was investigated. The cluster-encoded SARP B-004_07379 shared 38% amino acid identity and 51% similarity with PapR2.

To test the effect of *papR2* overexpression on naphthocyclinone biosynthesis, *Streptomyces* sp. B-004^T^ was transformed with the integrative plasmid pDS300 (12), in which *papR2* transcription is driven by the constitutive *ermE\** promoter. The resulting overexpression strain *Streptomyces* sp. B-004^T^/pDS300 and the control strain *Streptomyces* sp. B-004^T^/pIJ10257 carrying the empty vector (39) were cultivated in R5 medium for six days. Culture samples were collected every 24 h from days 2 to 6 of fermentation, extracted with ethyl acetate, and analyzed with LC-MS.

LC-MS analysis revealed that the *papR2* overexpression strain accumulated substantially higher amounts of naphthocyclinones than the strain with the empty-vector control as early as day 2 of cultivation (Fig. 5A). This biosynthetic difference became even more pronounced after three days of fermentation, with the overexpression strain producing 2.5-fold more α-naphthocyclinone and accumulating significant amounts of β-naphthocyclinone-epoxide compared to the control. Although naphthocyclinone production increased in both strains during prolonged cultivation, the relative difference between the overexpression strain and the control gradually diminished. The maximum production was observed after four days of growth for *Streptomyces* sp. B-004^T^/pDS300, whereas the control strain reached its peak after five days of cultivation (Fig. 5A).

**Fig. 5.**
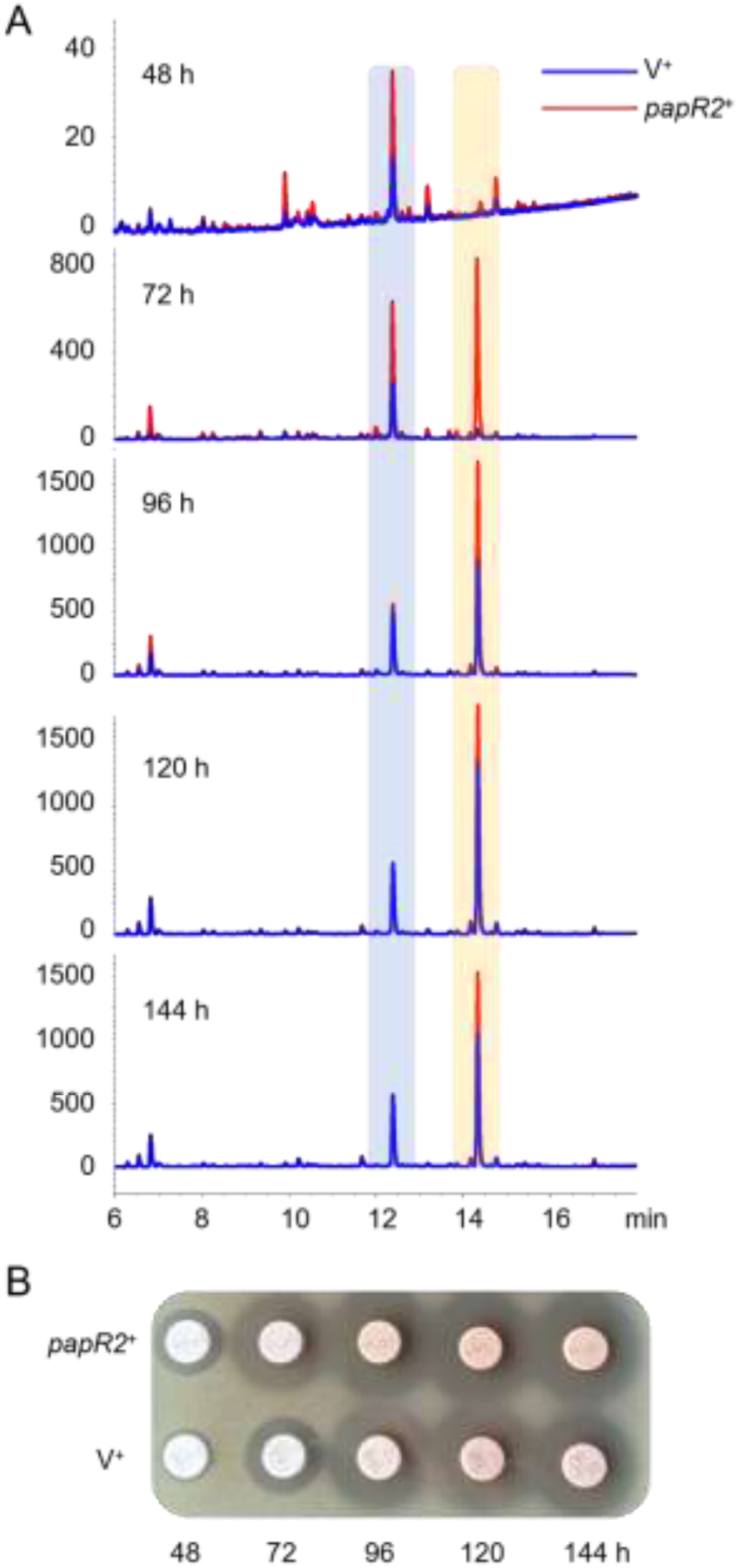
Comparative time-course analysis of naphthocyclinone production and antimicrobial activity in *Streptomyces* sp. B-004^T^ and its *papR2*⁺ overexpression strain. (**A**) Overlaid HPLC-UV chromatograms (254 nm) tracking the metabolic kinetics of extracts sampled at consecutive fermentation time points (48, 72, 96, 120, and 144 h). Blue traces represent the wild-type strain carrying the empty vector; red traces are the *papR2* overexpression mutant. Shaded vertical zones highlight the elution windows of the major accumulated naphthocyclinone congeners, α-naphthocyclinone (blue shaded) and β-naphthocyclinone-epoxide (yellow shaded). (**B**) Disc diffusion assay against *K. rhizophila* with the corresponding extracts used for HPLC analysis in panel **A**.

In parallel, the extracts were evaluated for antibacterial activity against *K. rhizophila.* Consistent with the LC-MS data, extracts from the *papR2* overexpression strain showed significantly larger inhibition zones against *K. rhizophila* than extracts from the control strain. This enhanced antibacterial effect was most pronounced for extracts obtained after 2 and 3 days of cultivation, directly correlating with the increased naphthocyclinone accumulation (Fig. 5B). These findings demonstrate that increased naphthocyclinone production directly translates into enhanced antibacterial activity. Furthermore, these data show that SARPs act as functional activators of the naphthocyclinone BGC.

### Naphthocyclinone production is triggered by co-isolated sympatric bacteria

Interestingly, during subsequent laboratory subculturing of *Streptomyces* sp. B-004^T^ on solid media, we noticed a distinct phenotypic instability; the strain gradually lost its ability to produce the characteristic reddish-brown pigmentation associated with naphthocyclinone production, yielding uncolored colonies after multiple generations. This morphological change was consistently correlated with a reduction in antibacterial activity. To systematically investigate this phenomenon, we monitored the strain under routine laboratory cultivation. The original cryopreserved stock of *Streptomyces* sp. B-004^T^ was repeatedly subcultured over 20 consecutive passages on solid SFM medium, with a cultivation time of 7 days for each round. Thereafter, naphthocyclinone production was completely abolished and its characteristic reddish-brown pigmentation had disappeared (Fig. 6A).

**Fig. 6.**
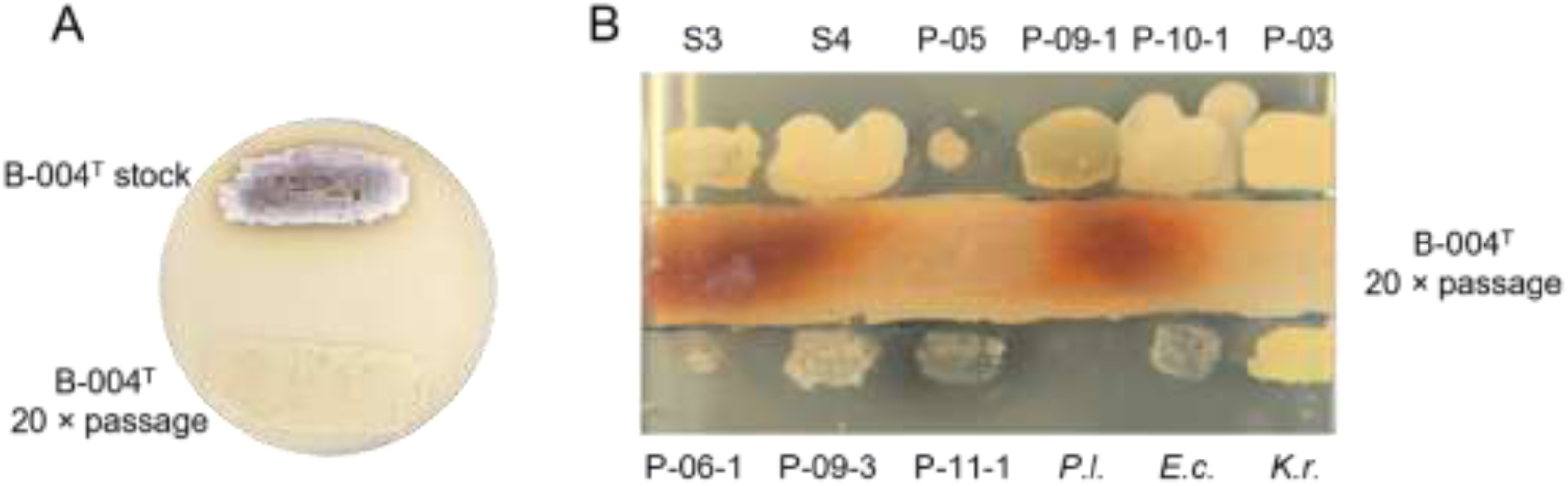
Phenotypic instability of naphthocyclinone production and interspecies metabolic cross-talk re-induces antibiotic biosynthesis in *Streptomyces* sp. B-004^T^. (**A**) Morphology of the *Streptomyces* sp. B-004^T^ strains grown on SFM medium for 5 days. The original cryopreserved stock (B-004 stock) and the strain obtained after 20 consecutive serial passages (B-004 20 × passage) show distinct pigmentation associated with production of reddish brown naphthocyclinone. (**B**) Co-cultivation assay demonstrating restoration of naphthocyclinone biosynthesis. The horizontal central line represents the non-producing *Streptomyces* sp. B-004^T^ strain (20 × passage) grown on R5A for 5 days. Vertical streaks correspond to sympatric nest co-isolates and reference strains (top row, left to right: *Bacillus* sp. S3, *Bacillus* sp. S4, *Peribacillus* sp. P-05, *Bacillus* sp. P-09-1, *Bacillus* sp. P-10-1, and *Priestia* sp. P-03; bottom row, left to right: *Paenibacillus* sp. P-06-1, *Metabacillus* sp. P-09-3, *Paenibacillus* sp. P-11-1, *Paenibacillus larvae* DSM 7030, *E. coli* Δ*tolC*, and *K. rhizophila* DSM 348). Localized accumulation of the dark reddish-brown naphthocyclinone pigment is selectively restored within the colony boundaries facing specific competitors.

Importantly, this loss of naphthocyclinone production was reversed during co-cultivation experiments. When the non-producing, repeatedly subcultured strain of *Streptomyces* sp. B-004^T^ was inoculated in the middle of an R5 agar plate, flanked by co-isolated *Bacilli*, naphthocyclinone production was re-induced in the zones facing some of the competitors (Fig. 6B). This re-induction indicated that the naphthocyclinone BGC remained functional but required specific interspecies signals for its activation.

### *Streptomyces* sp. B-004T represents a new species for which the name *Streptomyces hederae* sp. nov. is proposed

Morphological, biochemical, enzymatic, and chemotaxonomic properties of strain B-004^T^ were consistent with those of the genus *Streptomyces* (46). The strain exhibited good growth on ISP1, ISP2, ISP6, ISP7, Czapek’s agar, and NZ-amine media, whereas moderate growth was observed on ISP 3, ISP4, ISP5, and TSA agar media. Aerial mycelium developed only on ISP3 (cream-white, RAL 9001), ISP4 (green-beige with yellowish tinge, RAL 1000), and Czapek’s agar (grayish-white RAL 9002) after 10 days of incubation at 28°C. Optimal growth was observed at 25°-28°C and pH 6.5-7.5 in the absence of NaCl. The strain metabolized L-arabitol, D-adonitol, D-fucose, erythritol, D-lactose, D-melezitose, L-sorbose, and D-turanose, unlike its close relative *S. hokutonensis* DSM 102214^T^, which metabolized L-arabinose and methyl-a-D-glucopyranoside (Table S3). Both strains showed similar enzymatic profiles (Table S3). Strains B-004^T^ and DSM 102214^T^ contained *LL*-diaminopimelic acid in their peptidoglycan and had galactose, glucose, mannose, ribose, and rhamnose as whole-cell sugars. The major fatty acids (>10%) of strain B-004^T^ were *anteiso*-C15:0, *iso*-C16:0, and C16:0, whereas its close relative DSM 102214^T^ additionally contained *iso*-C14:0 (Table S3). The predominant menaquinones (>10%) of strain B-004^T^ were MK-9(H6) (23.3%), and MK-9(H8) (69.8%), whereas DSM 102214^T^ additionally contained MK-9(H4) (10.6%).

Pairwise 16S rRNA gene sequence similarity values between strain B-004^T^ and the type strains of validly named *Streptomyces* species were 99.7% with *Streptomyces prunicolor* NBRC 13075^T^ and *Streptomyces hokutonensis* R1-NS-1^T^, and 98.8% with *Streptomyces graminifolii* JL-22^T^ and *Streptomyces resistomycificus* NRRL ISP 5133^T^ (Fig. S6). These high 16S rRNA gene sequence similarities were consistent with the phylogenetic position of strain B-004^T^ in the nighbor-joining tree, in which it formed a well-supported subclade with *S. hokutonensis* R1-NS-1^T^ adjacent to *S. prunicolor* NBRC 13075^T^. These three strains were closely related to *S. graminifolii* JL-22^T^, together forming a distinct cluster (Fig. S6). Given the high 16S rRNA gene sequence similarity between strain B-004^T^ and its closest relative, exceeding the proposed threshold of 98.7% for prokaryotic species delineation, and the limited resolution of the 16S rRNA gene sequence for discriminating closely related *Streptomyces* species, genome-based taxogenomic analyses were performed.

Genome BLAST Distance Phylogeny (GBDP) showed that strain B-004^T^ formed a well-supported subclade with *S. hokutonensis,* adjacent to the type strains of *S. graminifolii and S. prunicolor* (Fig. 7). The dDDH values between strain B-004^T^ and the type strains of *S. hokutonensis*, *S. prunicolor, and S. graminifolii* were 56.4%, 40.9%, and 39.0 %, respectively, all below the 70% threshold for prokaryotic species delineation (47). These results were consistent with the corresponding ANIm values of 94.6% with *S. hokutonensis*, 91.1% with *S. prunicolor*, and 90.5% with *S. graminifolii*, which were below the generally accepted cutoff point of 95-96.7% for species delineation (48–52).

**Fig. 7.**
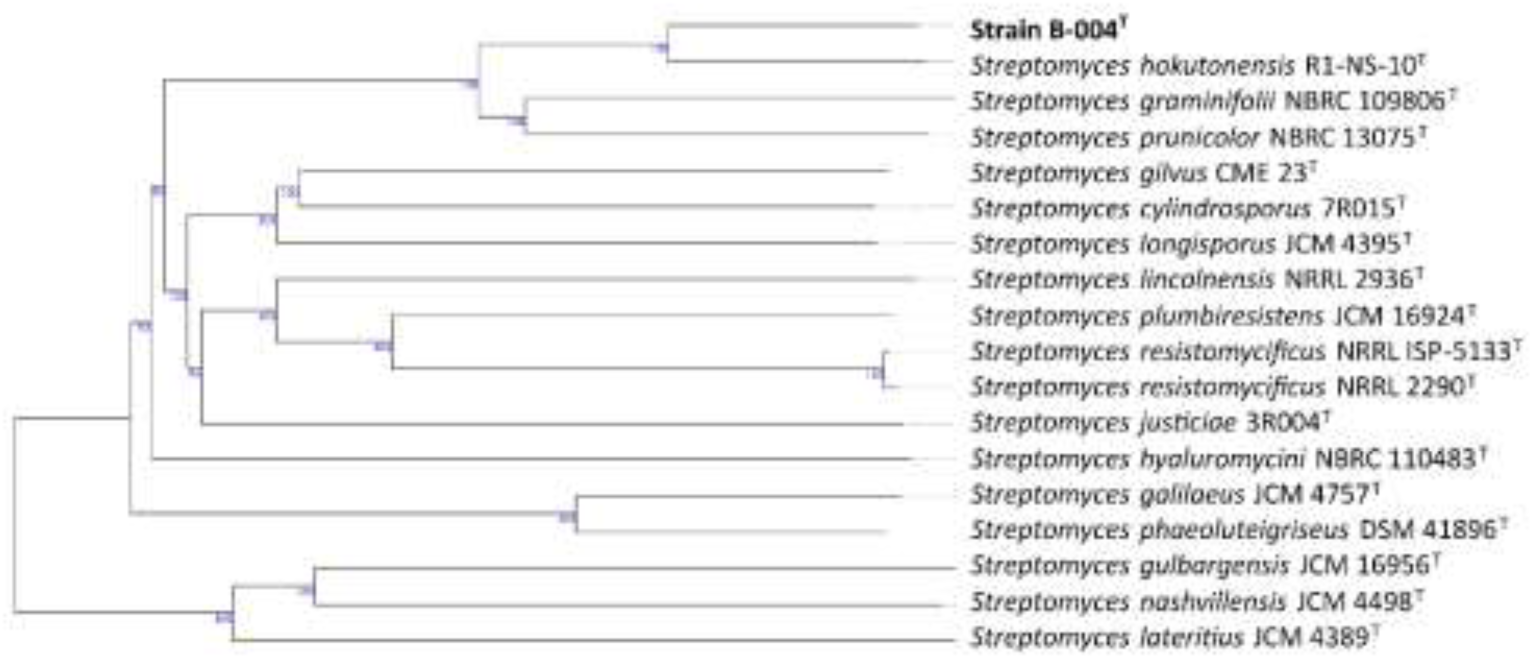
Genome BLAST Distance Phylogeny (GBDP) showing the phylogenetic relationship of strain B-004^T^ within the evolutionary radiation of the genus *Streptomyces*. Numbers above branches are GBDP pseudo-bootstrap support values >60 % from 100 replications. The tree was rooted at the midpoint.

Based on the phenotypic, chemotaxonomic, phylogenetic, and taxogenomic characteristics, strain B-004^T^ is proposed as the type strain of a novel species, for which the name *Streptomyces hederae* sp. nov. is proposed.

### Description of *Streptomyces hederae* sp. nov

*Streptomyces hederae* (he.de.ra.e, N.L. gen. fem. n. *hederae*, referring to *Colletes hederae*, the insect host with which the type strain was associated).

Aerobic, Gram-positive, filamentous actinobacterium with an extensively branched substrate mycelium. Aerial mycelium develops after 10 days of incubation at 28 °C and exhibits different colours depending on the growth medium: cream-white on ISP3, green-beige with a yellowish tinge on ISP4, and greyish-white on Czapek’s agar. Good growth occurs on ISP1, ISP2, ISP6, ISP7, Czapek’s agar, and NZ-amine media, whereas moderate growth is observed on ISP3, ISP4, ISP5, and TSA agar. Optimal growth occurs at 25–28 °C and pH 6.5–7.5 in the absence of NaCl. The peptidoglycan contains *LL*-diaminopimelic acid, and the whole-cell sugars comprise galactose, glucose, mannose, ribose, and rhamnose. The major fatty acids (>10%) are *anteiso*-C15:0, *iso*-C16:0, and C16:0. The predominant menaquinones (>10%) are MK-9(H6) (23.3%), and MK-9(H8) (69.8%).

The type strain B-004^T^ (= DSM 121525^T^ = LMG 34598^T^) was isolated from the fermented nest provision of a wild bee *Colletes hederae* collected in Braunschweig, Germany. The genome size is 11.19 Mb with G+C content of 70%. The Whole Genome Shotgun project has been deposited in DDBJ/ENA/GenBank under the accession number (ID:1522304, SUB16454956). The version described in this paper is version 1.0.

## Discussion

Although naphthocyclinones have been structurally and biosynthetically characterized for more than five decades, their ecological function has remained unknown. Our study provides the first experimental evidence that these aromatic polyketides mediate bacterial competition in an insect-associated microbial community. We identified a naphthocyclinone-producing *Streptomyces* strain B-004^T^ associated with the nest provisions of the solitary bee *C. hederae*, representing the first report of these metabolites in an insect-associated environment. Although underground nesting protects bees’ brood from abiotic fluctuations and predators, the high humidity and constant contact with soil makes provisions potentially prone to microbial spoilage (5). The risk of microbial contamination may be particularly high in nests of *C. hederae*, which forms dense nesting aggregations comprising thousands of individual nest cells in close proximity (53). Such high nest densities may further increase the spread of soil-associated microorganisms and the subsequent colonization of nest provisions. In this context, the presence of antibiotic-producing actinomycetes within the nests may contribute to the protection of the pollen provisions against microbial spoilage.

The antimicrobial activity observed for methanolic extracts from fermented nest provisions indicates that the antibacterial metabolites are present within the nests under natural conditions. The broad inhibitory activity of the *Streptomyces* sp. B-004^T^ supernatant against co-isolated sympatric *Bacilli* further supports an ecological role for naphthocyclinones in mediating microbial competition. Soil-associated bacteria, such as the genera *Bacillus*, *Peribacillus*, and *Paenibacillus*, were repeatedly isolated from nest provisions and may compete for nutrient resources within these nutrient-rich substrates. These microorganisms can originate either from soil or plants visited by the pollinator, making their complete exclusion from the collected material difficult. Notably, the strong inhibition of *P. larvae* by both the B-004^T^ wild-type culture supernatant and extract points to the potential role of naphthocyclinones in protecting nest provisions, while also suggesting a possible practical application. *P. larvae* is a spore-forming pathogen that causes American foulbrood, a disease capable of destroying honeybee colonies and threatening broader apiculture (7). Although *P. larvae* is primarily a honeybee pathogen, both wild solitary bees and managed honeybees frequently exploit the same floral resources and are therefore exposed to similar microbial communities, including potential pathogens (54). Beyond their ecological role in solitary bee nest protection, the potent activity of naturally occurring naphthocyclinones against *P. larvae* also highlights their potential as a source of antimicrobial leads for the control of American foulbrood. More broadly, this finding illustrates how ecologically informed screening can reveal the natural targets of specialized metabolites, which remain obscure in conventional antimicrobial screenings.

The present study expands our knowledge of the diversity of insect–actinomycete associations. Similar partnerships have been described for several insect lineages, including beewolves and fungus-growing ants, where microbial partners protect host resources or developing offspring against antagonistic microorganisms (14–16). Whether *Streptomyces* sp. B-004^T^ represents a stable symbiont of *C. hederae* or an environmentally acquired beneficial associate remains to be determined. While the majority of studies addressed the microbiome of honeybees and bumblebees, microorganisms associated with other solitary bee species remain poorly explored. Neither the distribution of actinobacteria across solitary bees nor the environmental sources of naphthocyclinone-producing *Streptomyces* are currently known. Resolving this question will be essential for understanding whether naphthocyclinone production represents a co-evolved defensive trait shaped by interactions between the host and its environment, or a more general ecological association. Ultimately, our findings highlight the potential of bee-associated environments as a rich source of actinomycetes producing ecologically relevant antibiotics.

Production of specialized metabolites in microorganisms comes at substantial costs. When propagated under favourable conditions, microorganisms may lose their ability to produce certain metabolites, undergo changes in growth properties, and alter their morphogenesis. In the present study *Streptomyces* sp. B-004^T^ gradually lost its capacity to synthesize naphthocyclinones upon repeated subculturing and long-term maintenance as a pure laboratory culture. Prolonged laboratory cultivation is frequently accompanied by a decline in secondary metabolite production. In some actinomycetes, this phenomenon has been associated with chromosomal rearrangements arising during repeated subculturing, which generate genetically heterogeneous populations of cells characterized by division of labor (55,56). Such rearrangements may be favored because they restrict the metabolic costs of specialized metabolite production to only a subset of cells, while the remaining cells benefit from the resulting division of labor. In addition, the loss of metabolite production may reflect the absence of naturally occurring environmental signals that arise through interactions with competitors or other members of the microbial community. Under laboratory conditions, the lack of such stimuli may lead to the downregulation or silencing of biosynthetic pathways, potentially reducing the metabolic costs associated with metabolite production when microbial competitors are absent. This hypothesis is also supported by our co-culturing experiments, showing that naphthocyclinone biosynthesis was restored in the repeatedly subcultured non-producing B-004^T^ strain during co-cultivation with *Bacilli* co-isolates. This suggests that the loss of production reflects environmentally regulated silencing rather than irreversible genetic mutations. The restoration of naphthocyclinone production during co-cultivation with co-isolates further confirms that its biosynthesis is sustained by ecological interactions within the native microbial community. In actinomycetes, cross-species induction of specialized metabolism is a well-recognized phenomenon, triggered by factors such as cellular debris, iron competition, or specific signaling molecules secreted by neighboring bacteria (57). Our findings add another ecological link showing that interactions within the *C. hederae* native nest microbiome provide signals required to maintain naphthocyclinone biosynthesis under natural conditions. Activating antibiotic production only in periods of active microbial interactions would minimize the metabolic costs associated with specialized metabolite biosynthesis, while preserving the ability to respond rapidly when competitors are present. More broadly, this observation provides a possible explanation for why many BGCs remain silent under standard laboratory conditions, where the ecological interactions that normally activate their expression are absent. Furthermore, it highlights why activation strategies, such as the expression of direct pathway-specific transcriptional regulators (e.g. SARP-type regulators) are particularly effective because they overcome the requirement for complex ecological activation signals, thereby enabling the targeted activation of otherwise silent biosynthetic pathways.

Together, our study links two dimensions of microbial natural product ecology that are often investigated separately: ecological function and interaction-dependent regulation. By starting from naturally co-occurring microorganisms in a spatially constrained bee-associated microhabitat, we connect a *Streptomyces* producer and its metabolites to susceptible natural targets and show experimentally that interactions with community members can regulate metabolite production. At the same time, the pronounced activity against *P. larvae* provides an additional dimension of biological relevance, identifying a long-known natural product as strongly active against a major bee pathogen. More broadly, our findings suggest that the apparent silence of specialized metabolite pathways under standard laboratory conditions, a phenomenon common among actinomycetes, may often reflect the absence of ecological signals involved in their regulation in native communities. By linking ecological function with biosynthetic regulation, this study provides insight into two long-standing questions in natural product research: what ecological roles specialized metabolites fulfill in nature, and why their biosynthetic pathways are so frequently silent despite remaining genetically intact.

## DATA AVAILABILITY

All data used in this study are included in the article and its Supporting Information files. The genome assembly of *Streptomyces* sp. strain B-004 (DSM 121525) has been submitted to the National Center for Biotechnology Information (NCBI) under submission number SUB16454956. The corresponding accession numbers for the genome assembly, GenBank assembly, NCBI RefSeq assembly, whole-genome shotgun (WGS) project, and BioSample will be provided upon acceptance of the manuscript. The BioProject accession number is PRJNA1522304.

## Contributions

RM conceived the study. RM and YM designed the research and developed the methodology. RM, IN, MJ, YK, AY, JS, CS, and BB conducted experiments. RM and IN prepared the visualizations. RM and YM drafted the initial manuscript. RM and YM supervised the work. All authors reviewed, edited and approved the final version of the manuscript.

## Acknowledgments

We thank Rüdiger Pukall for providing strain *Paenibacillus larvae* DSM 7030^T^ and Rüdiger Pukall, Sabine Gronow, Andrey Yurkov, and Imen Nouioui for depositing strains in the DSMZ collection. We gratefully acknowledge Sarah Kirstein and Meina Neumann-Schaal for fatty acid and quinone analyses and providing access to the HRMS instrumentation. We thank Gabriele Pötter and Meike Döppner for excellent technical assistance. We would like to thank Mark Buttner for providing pIJ10257. We are also grateful to Andriy Luzhetskyy for providing the pINT and pKGLP2 plasmids.

